# Knockout of Perilipin-2 in Microglia Alters Lipid Droplet Accumulation and Response to Alzheimer’s Disease Stimuli

**DOI:** 10.1101/2025.09.02.673815

**Authors:** Isaiah O. Stephens, Lance Johnson

## Abstract

Lipid droplets (LDs) are emerging as critical regulators of cellular metabolism and inflammation, with their accumulation in microglia linked to aging and neurodegeneration. Perilipin 2 (Plin2) is a ubiquitously expressed LD-associated protein that stabilizes lipid stores, and in peripheral tissues its upregulation promotes lipid retention, inflammation, and metabolic dysfunction. However, the role of Plin2 in brain-resident microglia remains undefined. Here, we used CRISPR-engineered Plin2 knockout (KO) BV2 microglia to investigate the contribution of Plin2 to lipid accumulation, bioenergetics, and immune function. Compared to wild-type (WT) cells, Plin2 KO microglia exhibited markedly reduced LD burden under both basal and oleic acid–loaded conditions. Functionally, this was associated with enhanced phagocytosis of zymosan particles, even after lipid loading, indicating improved clearance capacity in the absence of Plin2. Transcriptomic analyses revealed genotype-specific responses to amyloid-β (Aβ), particularly in pathways related to mitochondrial metabolism. Seahorse assays confirmed that Plin2 KO cells adopt a distinct bioenergetic profile, with reduced basal respiration and glycolysis but preserved mitochondrial capacity, increased spare respiratory reserve, and a blunted glycolytic response to Aβ. Together, these findings identify Plin2 as a regulator of microglial lipid storage and metabolic state, with its loss alleviating lipid accumulation, improving phagocytic function, and altering Aβ-induced metabolic reprogramming. Targeting Plin2 may therefore represent a potential strategy to modulate microglial metabolism and function in aging and neurodegeneration.

## INTRODUCTION

Lipid droplets (LDs) are dynamic intracellular organelles that serve as neutral lipid storage depots and play central roles in lipid metabolism. Structurally, LDs consist of a neutral lipid core surrounded by a phospholipid monolayer embedded with proteins that regulate lipid storage, mobilization, and organelle interactions^1,2^. They form from the endoplasmic reticulum and interact extensively with mitochondria, lysosomes, and peroxisomes, coordinating metabolic flux^1,3^. LDs respond to cellular energy demands by storing excess fatty acids or releasing them via lipolysis and lipophagy^1,4^. They buffer toxic lipid species and support immune signaling by serving as platforms for eicosanoid synthesis, including prostaglandins and leukotrienes^5-7^. These immunometabolic roles implicate LDs in a diverse range of diseases.

LD accumulation is a feature of many chronic inflammatory and metabolic diseases, including NAFLD, atherosclerosis, obesity, and type 2 diabetes, where it contributes to inflammation and insulin resistance^8-11^. Although LD research has historically focused on peripheral tissues, growing evidence highlights their significance in the central nervous system (CNS), particularly within glial cells like astrocytes and microglia^12,13^. In microglia, LDs form in response to aging, metabolic stress, inflammation, or uptake of lipid-rich debris such as myelin^14^. Under these conditions, microglia can transition into a lipid-laden phenotype known as lipid droplet–accumulating microglia (LDAM), characterized by elevated LD content, impaired phagocytosis, and a pro-inflammatory transcriptional profile^12,15^. LDAM are induced by stimuli such as lipopolysaccharide (LPS) and amyloid-beta (Aβ), which also elevate ROS and cytokine production while impairing microglial clearance functions^12,16,17^. These cells are increasingly observed with age and are enriched near amyloid plaques in Alzheimer’s disease (AD), where their abundance correlates with disease progression^18^.

Plin2, a lipid droplet–associated protein, regulates lipid storage and mobilization by stabilizing droplets and limiting lipase access^19,20^. In peripheral tissues, Plin2 upregulation promotes lipid retention in settings such as fatty liver, atherosclerosis, and insulin resistance^9,21,22^, while Plin2 deficiency reduces lipid burden and inflammation^23,24^. In the CNS, Plin2 is often used as an LD marker, with expression increasing with age, stroke, and AD^12,15,18^, where it is prominently detected in lipid-laden microglia. However, its precise function in glial lipid biology, particularly in neurodegeneration, remains unclear, representing a critical gap with important implications for neuroinflammation and disease.

Given Plin2’s central role in lipid droplet biology and its elevated expression in aging and disease-associated microglia, we sought to define its functional contribution to microglial responses. To do so, we created a CRISPR-generated Plin2 knockout BV2 microglial model and compared it to wild-type cells under multiple AD-relevant stimulatory conditions. We demonstrate that loss of Plin2 reduces lipid burden, modulates inflammatory signaling, and restores phagocytic function in microglia. These findings identify Plin2 as a potential therapeutic target for modulating neuroinflammation in aging and neurodegeneration.

## METHODS

### BV2 Cell Culture

CRISPR/Cas9-generated Plin2 knockout (Plin2 KO) BV2 microglial cells and their wild-type (WT) counterparts were generated with Ubigene (Guangzhou, China). Cells were maintained in DMEM/F-12 supplemented with 1× GlutaMAX, 10% fetal bovine serum (FBS), and 1% penicillin-streptomycin. Cultures were incubated at 37 °C in a humidified atmosphere containing 5% CO2 under normoxic conditions. For experiments, cells were seeded onto Poly-L-Lysine (PLL)-coated culture vessels (6-well plates or 8-well chamber slides). PLL was applied for 2 hours at 37 °C, followed by a 1× DPBS wash and air-drying overnight.

For omics-based experiments, BV2 cells were treated with Alzheimer’s disease-relevant stimuli for 24 hours prior to RNA or lipid extraction. Treatments included oleic acid (250 μM), myelin debris (15 μg/cm^2^), differentiated N2a neuron-like cells (dN2A) at a 5:1 dN2A:BV2 ratio, and amyloid-β (1.5 μM).

Oleic acid was purchased pre-conjugated to bovine serum albumin (BSA) at 100mg/ml in DPBS (Sigma Cat: 03008 Lot: SLCH8543). Stock concentration reported as 892 μg/ml, equivalent to a 3.14mM concentration. This stock was spiked into an aliquot of growth media to a concentration of 250 μM on day of treatment. Oleic acid containing media was then used during media change for experimental wells/chambers. Control wells received media spiked with an equivalent volume of DPBS containing only 100mg/ml BSA.

Synthetic amyloid-β (1-42) (AnaSpec, San Jose, CA) prepared as previously described^25^. A 1mg stock of lyophilized amyloid-β was dissolved in 200 μl of hexafluoroisopropanol, divvied out into 0.2mg aliquots, dried in a chemical fume hood, and then stored as a film at −20 °C. Immediately prior to use, amyloid-β was resuspended in Me_2_SO to 10 mM and water bath sonicated for 10 min. It was further prepared by diluting the amyloid-β to 25 μM with phenol red-free DMEM-F12 and incubating for 24 hours at 4 °C without shaking.

N2A cells were split into a few dozen T-75 flasks with non-filter cap lids. Once cells grew to confluency, the caps were tightened to induce hypoxia and cell death. Cells began to detach from the flasks and all apoptotic cells and media were collected, split into 8 × 10^6^ cells/ml aliquots and frozen for later experimental use.

Myelin debris was isolated from adult mouse brain using Percoll density gradient centrifugation as previously described^26^. Briefly, brains were homogenized in ice-cold buffer and layered onto a Percoll gradient, then centrifuged to separate myelin-rich fractions. The upper myelin-containing layer was collected, pelleted by centrifugation, and washed twice in sterile 1× DPBS by resuspension and re-pelleting. The final pellet was resuspended in 1× DPBS and homogenized by repeated passage through a fine-gauge needle to ensure uniform dispersion. Aliquots were then stored at −80 °C until time for experimental use.

### Mouse Model

Human APOE4 targeted-replacement (APOE4-TR) mice were crossed with 5xFAD transgenic mice to generate APOE4-TR;5xFAD offspring which were aged to 12 months and used for plaque associated Plin2 microglia validation. Animals were housed on a 12:12-h light/dark cycle with food and water provided ad libitum. All animal work was performed in accordance with institutional guidelines and approved by the University of Kentucky IACUC (2016-2569).

### Lipid Droplet Staining

For lipid droplet (LD) visualization, BV2 cells were treated with 250 μM oleic acid (OA) or 1xDPBS as a control for 24 hours in glass chamber slides. After treatment, cells were washed with 1× DPBS and incubated in OptiMEM containing 0.15 μM LipiGreen (Dojindo) for 30 minutes at 37 °C. Cells were then fixed in 4% paraformaldehyde (PFA) for 15 minutes, washed three times with 1× DPBS, and mounted using Vectashield HardSet Antifade Mounting Medium with DAPI. Slides were cured overnight and imaged using a Nikon AXR confocal microscope (University of Kentucky Light Microscopy Core).

### LD Time Course Experiment

BV2 microglial cells were seeded on glass chamber slides in standard growth medium and allowed to adhere overnight at 37 °C/5% CO2. The following morning, cells were swapped to media with either 250 µM oleic acid or 1.5 µM amyloid-β in growth medium. Slides were removed at 6, 18, and 24 h post-treatment and washed once with 1× DPBS before incubation in OptiMEM containing 0.15 µM LipiGreen for 30 min at 37 °C. Cells were then fixed in 4% paraformaldehyde for 15 min, washed three times with 1x DPBS, and mounted in Vectashield HardSet with DAPI. Immediately after the 24 hour synthesis time point, remaining slides were washed and switched to serum-free media to promote LD degradation; these were processed at +2, +6, +18, and +24 hours after serum removal using the identical LipiGreen staining, fixation, and mounting steps. All images were acquired on a Nikon AXR confocal microscope (University of Kentucky Light Microscopy Core).

### Phagocytosis Assay

Following 24-hour treatment with or without OA, cells in 8-chamber slides were incubated with either 15 μL of 1× DPBS (vehicle) or 15 μL of pHrodo™ Red Zymosan Bioparticles (Abcam) per well. After a 2-hour incubation at 37 °C, cells were washed and fixed with 4% PFA for 15 minutes, then mounted using Vectashield hard-set mounting media with DAPI and allowed to cure overnight hidden from light. Slide were then prepared and imaged on the Nikon W1 spinning disk confocal microscope using a 40x water immersion objective (University of Kentucky Light Microscopy Core).

### Lipidomics

BV2 cells were plated on poly-l-lysine-coated 6-well plates and grown to ∼75% confluence at 37 °C, 5% CO_2_. On the day of extraction, plates were removed from the incubator and placed in the biosafety cabinet. Medium was aspirated, wells were rinsed once with 500 µL ice-cold 1× DPBS and aspirated again, then 500 µL of pre-chilled extraction solvent (50:50 methanol:butanol, 10 mM ammonium formate) was added to each well. Plates were transferred to –80 °C for 15 minutes, then cells were scraped in that first 500 µL and transferred to 1.5 mL tubes. A second 500 µL solvent wash was used to rinse each well; that wash was pooled into the same tube (≈1 mL total). Tubes were vortexed briefly and stored at –80 °C until the day of analysis. On analysis day, samples were thawed on ice, centrifuged at 14,000 × g for 10 minutes at 4 °C to pellet debris, and the clear supernatant was transferred to LC-MS vials. Lipidomic data were acquired on an Agilent QQQ LC-MS system.

### Transcriptomics

Total RNA was isolated from BV2 cells using the Qiagen RNeasy Mini Kit. RNA quality was assessed, and samples with RIN ≥ 8 were submitted to Novogene for poly(A)-enriched paired-end (2×150 bp) Illumina sequencing. Reads were aligned to the mouse reference genome (GRCm38/mm10), and gene counts were obtained. Differential gene expression was analyzed in R using the DESeq2 package. Genes with FDR adjusted p-values < 0.05 and fold change ≥ 1.5 were considered differentially expressed. Principal component analysis (PCA), hierarchical clustering, and Gene Ontology (GO) enrichment analyses were performed using prcomp, pheatmap, and clusterProfiler packages, respectively. All data can be found in Supplementary Table 1.

### Co-expression network construction

BV2 bulk RNA-seq counts and metadata were loaded into R. Genes with CPM > 1 in ≥ 25% of samples, zero-variance genes, and Rps/Rpl transcripts were removed. Raw counts were variance-stabilized with DESeq2’s vst (dds, blind=TRUE), outliers were removed by hierarchical clustering, and the cleaned matrix was used for WGCNA module detection. A signed co-expression network was built in WGCNA (v1.71) using soft-threshold power 6 (scale-free R^2^ ≥ 0.8), blockwiseModules (minModuleSize = 30, deepSplit=3, mergeCutHeight=0.20), and TOMType=“signed.” Module eigengenes were correlated with Genotype_Treatment groups (Pearson’s r; p-values from corPvalueStudent), and intramodular connectivity (kME) defined hub genes. Module gene sets were tested for GO Biological Process enrichment with clusterProfiler (BH-adj p < 0.05, q < 0.05, minGSSize = 5). Adjacency and node-attribute tables (module color and kME) for each module and for hub-gene subsets (top 10 ranked by positive kME) were exported via exportNetworkToCytoscape for visualization.

### Seahorse Oxidation Assay

BV2 cells (WT and Plin2 KO) were seeded at 8 × 10^3^ cells/well in Agilent XF96 plates and allowed to adhere overnight at 37 °C/5% CO_2_. Cells were then treated ± 1.5 µM amyloid-β for 24 h, washed twice with Seahorse XF DMEM assay medium (10 mM glucose, 2 mM l-glutamine, 1 mM sodium pyruvate, pH 7.4), and equilibrated at 37 °C (no CO_2_) for 45 min. OCR was measured on an XFe96 Analyzer using the XF Cell Mito Stress Test (1 µM oligomycin, 2 µM FCCP, 0.5 µM rotenone/antimycin A), with three 3 min mix–2 min wait–3 min measure cycles per phase (cycles 1–3 basal, 4–6 ATP-linked, 7–9 maximal, 10–12 non-mitochondrial). We calculated basal respiration, ATP production (basal – oligomycin), proton leak (oligo – non-mito), maximal respiration (FCCP – non-mito), spare respiratory capacity [(maximal–basal)/basal × 100], and coupling efficiency (ATP production/basal × 100). Each condition was run in up to 12 technical replicates (injector-failure wells excluded), and OCR values were normalized to DAPI-stained nuclear counts.

## RESULTS

### Plin2 knockout lowers lipid droplet accumulation and size in BV2 cells after oleic acid stimulation

To establish the role of Plin2 in microglial LD biology, we first examined its expression in vivo. Similar to previous reports^18,27^, immunofluorescence staining in 5xFAD mice revealed Plin2-positive microglia clustered around amyloid plaques, where they contained abundant LDs (Fig. 1a).

**Figure 1.**
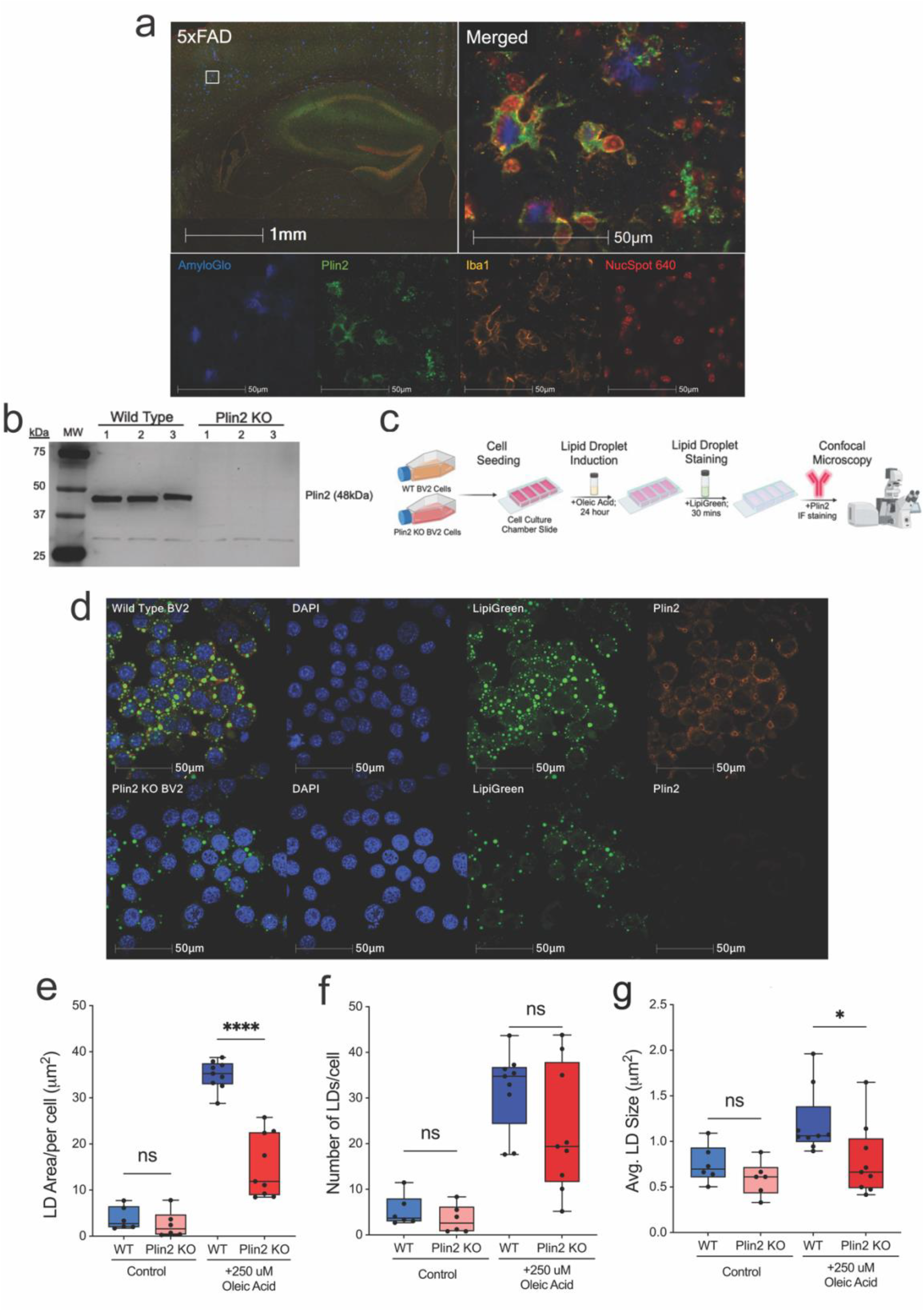
Plin2 knockout reduces oleic acid–induced lipid droplet accumulation in BV2 microglia. (a) Immunofluorescent staining of 5xFAD brain sections showing Plin2-positive (green) microglia (Iba1, yellow) enriched around amyloid plaques (AmyloGlo, blue) with lipid droplet accumulation (NucSpot, red). Inset highlights plaque-associated Plin2+ microglia. (b) Western blot validation of Plin2 knockout (KO) in BV2 cells using CRISPR-Cas9, confirming loss of the 48 kDa Plin2 protein. (c) Experimental workflow schematic for oleic acid (OA) loading, lipid droplet staining, and confocal imaging. (d) Representative confocal images of WT and Plin2 KO BV2 cells under basal and OA-loaded conditions, stained with LipiGreen for lipid droplets (green), Plin2 (red), and DAPI (blue). (e–g) Quantification of lipid droplet parameters in WT and KO cells. (e) Total lipid droplet area per cell (****p < 0.0001). (f) Number of lipid droplets per cell (ns). (g) Average lipid droplet size (*p < 0.05). Data are mean ± SEM; individual points represent the average of 3 separate images taken per well.

To directly test the functional contribution of Plin2, we first confirmed the loss of Plin2 in our KO microglia with the absence of the 48 kDa Plin2 protein confirmed by Western blotting (Fig. 1b). We then employed an oleic acid (OA)–loading paradigm to assess LD formation, as outlined in the experimental workflow schematic (Fig. 1c). Under basal conditions, both WT and Plin2 KO cells exhibited few LDs. Following 24 hours of OA treatment, WT cells displayed a robust increase in total LD area, whereas this accumulation was significantly attenuated in Plin2 KO cells (Fig. 1d,e; ****p < 0.0001). The number of LDs per cell was not significantly altered between genotypes (Fig. 1f), but the average droplet size was significantly reduced in KO cells compared to WT (Fig. 1g; *p < 0.05). Together, these findings demonstrate that Plin2 is required for efficient expansion of lipid droplet size and accumulation in microglia following an exogenous lipid challenge.

### Phagocytosis of zymosan particles is enhanced in Plin2 KO cell

Lipid-laden phagocytes, including LDAMs, have been shown to be deficient in their ability to clear various targets^12^. To determine if the loss of Plin2 and lower accumulation of LDs influences phagocytic capacity, we tested the ability of Plin2 KO cells to clear zymosan particles under basal and OA-loaded conditions. Zymosan, a yeast-derived particle that robustly engages pattern recognition receptors, is a standard phagocytosis assay and provides a reliable readout of microglial clearance function in this context^12,28,29^. Under control conditions, Plin2 KO cells internalized significantly more zymosan than WT (Fig. 2a–b; ****p < 0.0001). Treatment with 250 µM oleic acid for 24 hours had no effect on WT uptake (WT + OA vs. WT control: ns), yet further increased phagocytosis in KO cells (**p < 0.01). Even after lipid loading, KO cells exhibited substantially greater uptake than WT + OA (****p < 0.0001), demonstrating that Plin2 constrains phagocytic capacity in BV2 microglia under both normal and lipid-loaded conditions. Together, these results underscore the impact of lipid accumulation on phagocytic function and suggest that reducing Plin2 may augment microglial clearance capacity.

**Figure 2.**
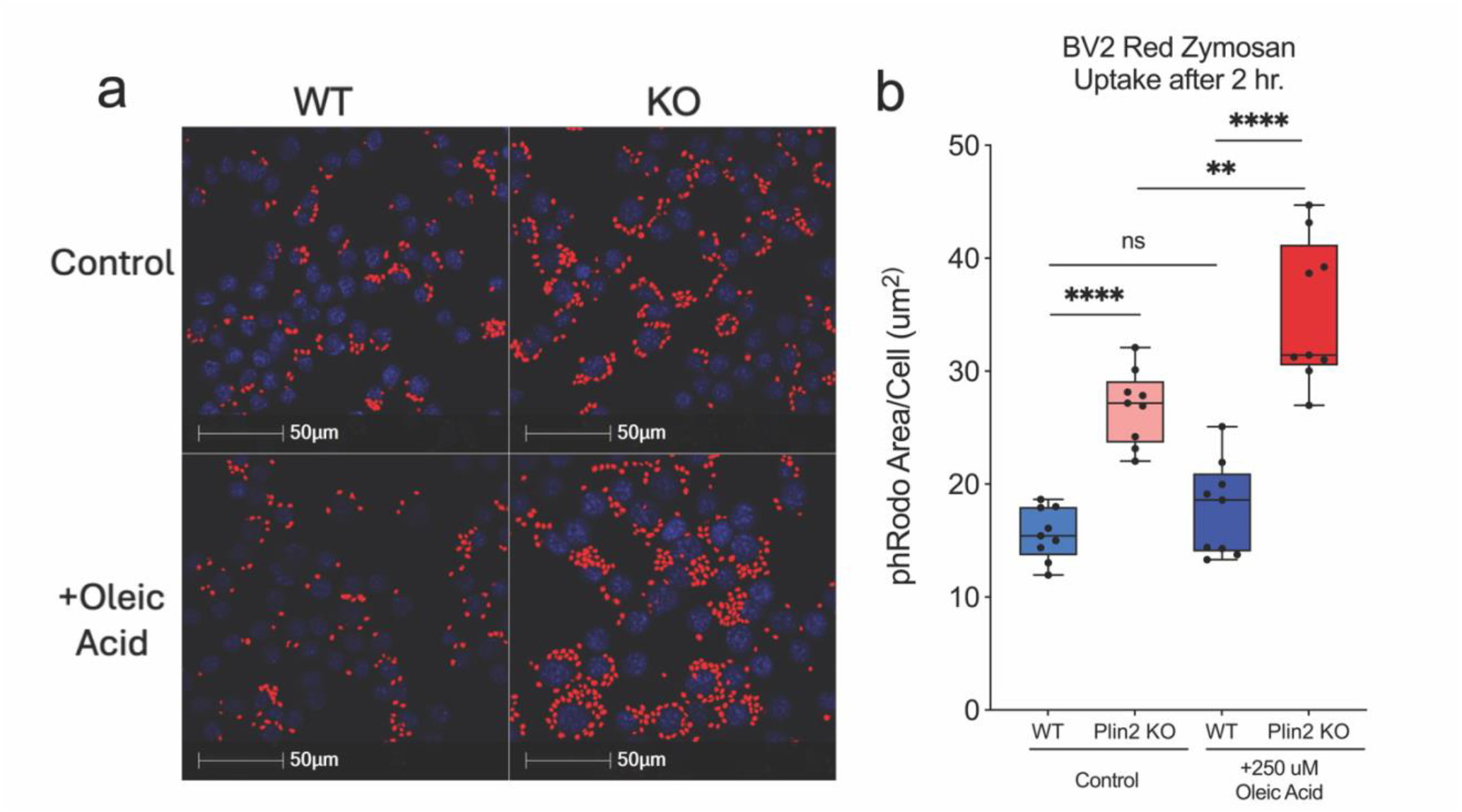
Oleic acid–dependent enhancement of zymosan uptake in Plin2-deficient microglia. BV2 cells (WT and Plin2 KO) were exposed for 24 h to vehicle or 250 µM oleic acid, then challenged with pHrodo Red Zymosan Bioparticles for 2 h. (a) Representative confocal images of internalized bioparticles (red) and DAPI-stained nuclei (blue). (b) Mean pHrodo-positive area per cell quantified from nine wells per condition (three fields averaged per well; total = 27 fields). Two-way ANOVA with Sidak’s correction: ****p < 0.0001 versus WT + vehicle; **p < 0.01 versus KO + vehicle; ns, WT + OA versus WT + vehicle.

### Bulk RNA-Sequencing Shows Plin2 Alters BV2 Response Profiles to AD-Relevant Stimuli

To investigate how loss of Plin2 reprograms microglial transcriptional programs, we performed bulk RNA-seq under basal and stimulated conditions and complemented this with Weighted Gene Co-expression Network Analysis (WGCNA) to resolve coordinated gene networks. Bulk RNA-seq revealed clear transcriptional divergence between Plin2 KO and WT microglia, with the extent of separation varying by treatment (Fig. 3a-b). At large, the KO cells showed upregulation of innate immune, chemotaxis, and migration pathways, while sterol and cholesterol biosynthesis and proliferative programs were downregulated (Fig. 3c, f– h). Importantly, this immune-versus-lipid metabolic split was evident not only at baseline but also under AD-relevant stimuli such as myelin and apoptotic neurons. By contrast, Aβ exposure elicited a distinct transcriptional response: KO cells upregulated mitochondrial and oxidative phosphorylation pathways, while small GTPase-mediated signaling and cytoskeletal regulation were downregulated (Fig. 3d–e).

**Figure 3.**
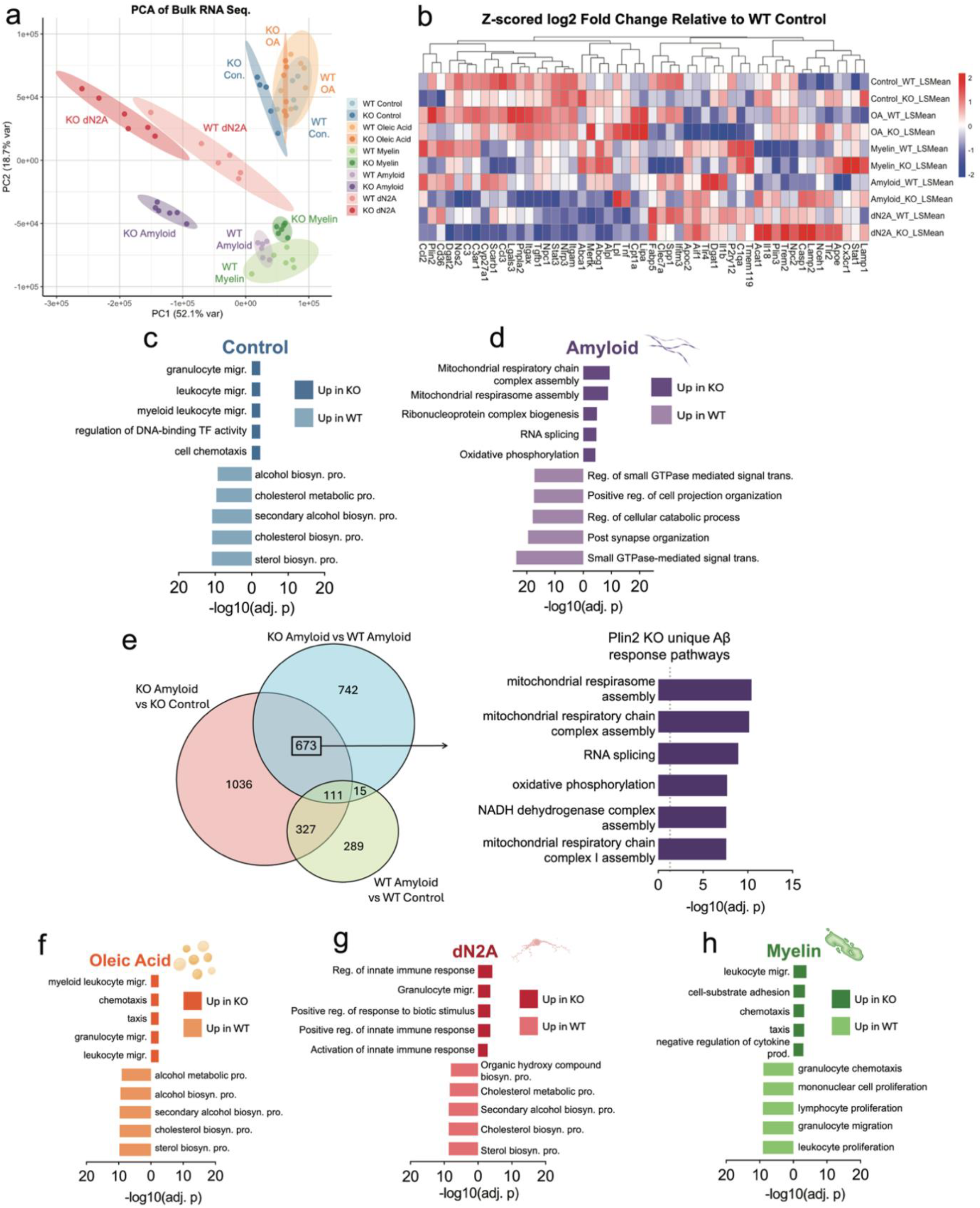
Plin2 deletion reprograms the microglia transcriptomes across stimuli. Principal component analysis (PCA) **(a)** of bulk RNA-seq from WT and Plin2 KO BV2 cells under control, amyloid beta (Aβ), oleic acid (OA), dN2A, and myelin treatments. **b**, Heat map of microglial genes and representative differentially expressed genes (DEGs), shown as Z-scored log2 fold change relative to WT Control (scale at right). **c–d, f–h**, Pathway enrichment analyses for DEGs in Control (**c**), Aβ (**d**), OA (**f**), dN2A (**g**), and Myelin (**h**). Bars show the top enriched Gene Ontology terms for genes up in KO (right bars) and up in WT (left bars), ranked by –log10(adjusted p value). Across conditions, WT preferentially enriches sterol and broader lipid-biosynthetic programs, whereas KO elevates innate immune, cytoskeletal, vesicle/lysosome, and (under Aβ) mitochondrial respiration/respirasome pathways. **e**, Venn diagram of DEGs under Aβ exposure across three contrasts: KO Aβ vs KO Control, WT Aβ vs WT Control, and KO Aβ vs WT Aβ.

WGCNA further resolved these networks, identifying KO-associated modules (cyan, pink, tan, black) that were consistently upregulated across all treatment conditions, and capturing programs related to Fc receptor signaling, innate immune activation, cytoskeletal remodeling, and cell cycle regulation (Fig. 4c–d). Notably, the cyan module contained Fcer1g, a central adaptor for Fc receptor signaling, while the tan module included DNA replication and cell cycle regulators such as Mcm3 and Cdc6 (Fig. 4d). Conversely, modules that were consistently downregulated in KO (purple, red, green-yellow, magenta) contained pathways linked to sterol and cholesterol biosynthesis, TNF-driven cytokine signaling, and adaptive immune activation (Fig. 4c). Key hub genes within these downregulated modules included canonical lipid regulators (Hmgcr, Hmgcs1) and immune mediators (Clec7a, Spp1) (Fig. 4e).

**Figure 4.**
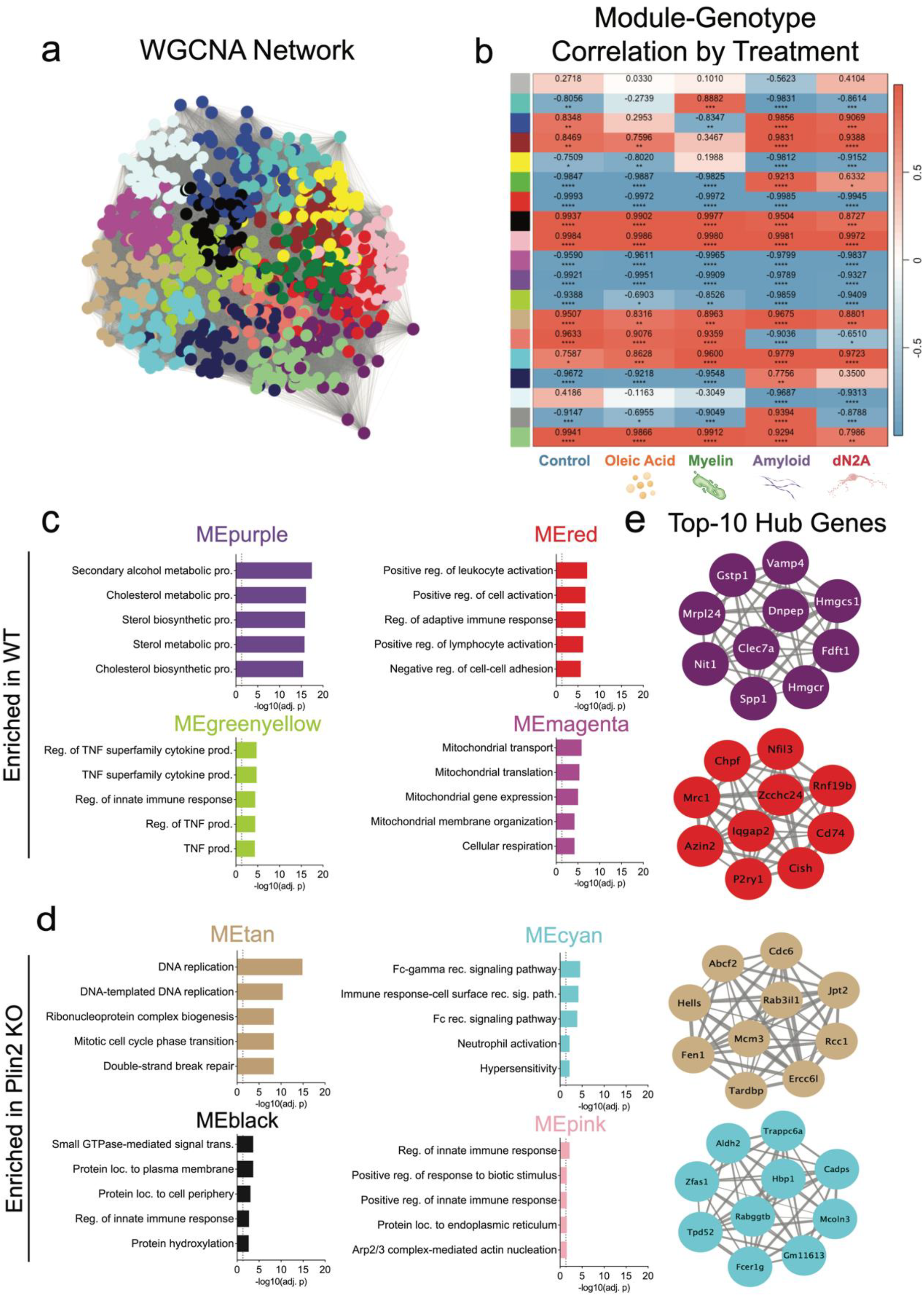
Co-expression network analysis links genotype to pathway modules and hub genes. **a**, WGCNA network built from variance-stabilized RNA-seq counts across all samples and treatments. Nodes are genes, edges reflect topological overlap, colors denote data-driven modules. **b**, Heat map of module eigengene correlation with genotype within each treatment. Values are Pearson r with correspondingadj. p values shown in each cell. Red indicates a positive correlation with the Plin2 KO, blue indicates a negative correlation. **c**, Modules enriched in WT. Bars show top Gene Ontology terms for representative WT-associated modules (for example MEpurple, MEred, MEgreenyellow, MEmagenta), ranked by –log10 of the adjusted p value. **d**, Modules enriched in Plin2 KO. Bars show top terms for representative KO-associated modules (for example MEtan, MEcyan, MEblack, MEpink), ranked by –log10 of the adjusted p value. **e**, Top-10 hub genes for selected modules. Networks display the ten genes with highest intramodular connectivity (kME) per module; edge thickness reflects connection strength within the module. Data on each WGCNA module can be found in WGCNA Supplemental Table.

Taken together, bulk RNA-seq and WGCNA demonstrate that Plin2 shapes both metabolic and immune programs in microglia. Across multiple stimuli, KO cells consistently downregulated sterol and lipid biosynthetic pathways while upregulating immune and Fc receptor–associated networks, with the strongest divergence observed under Aβ challenge. These transcriptomic differences provide a mechanistic framework for the altered lipid handling and phagocytic phenotypes observed in Plin2 KO microglia.

### Plin2 Regulates Lipid Droplet Dynamics in Response to Oleic Acid and Amyloid-β

As our transcriptomic analyses indicated that Aβ elicited the strongest genotype-specific transcriptional response in Plin2 KO versus WT microglia, we next examined whether these differences extended to LD dynamics. We performed a two-phase time course in our BV2 cells treated with 250 µM OA or 1.5 µM Aβ, measuring LD synthesis at 6, 18, and 24 hours, followed by a serum-free chase at +2, +6, +18, and +24 hours to monitor degradation.

Under OA, Plin2 KO cells accumulated significantly fewer LDs than WT at 18 hours (*p < 0.05) and 24 hours (**p < 0.01) (Fig. 5c). After serum removal, both genotypes showed progressive LD clearance, returning to baseline by +24 hours; however, KO cells maintained consistently lower LD levels throughout the time course. With Aβ, KO cells again showed no significant LD accumulation during the first 24 hours, whereas WT displayed a marked increase, detectable at 6 (*p < 0.05), peaking at 18 (****p < 0.0001), and still elevated at 24 hours (*p < 0.05) (Fig. 5d). Together, these findings show that loss of Plin2 limits LD accumulation in response to both OA and Aβ, underscoring its role in regulating microglial lipid storage in response to diverse stimuli.

**Figure 5.**
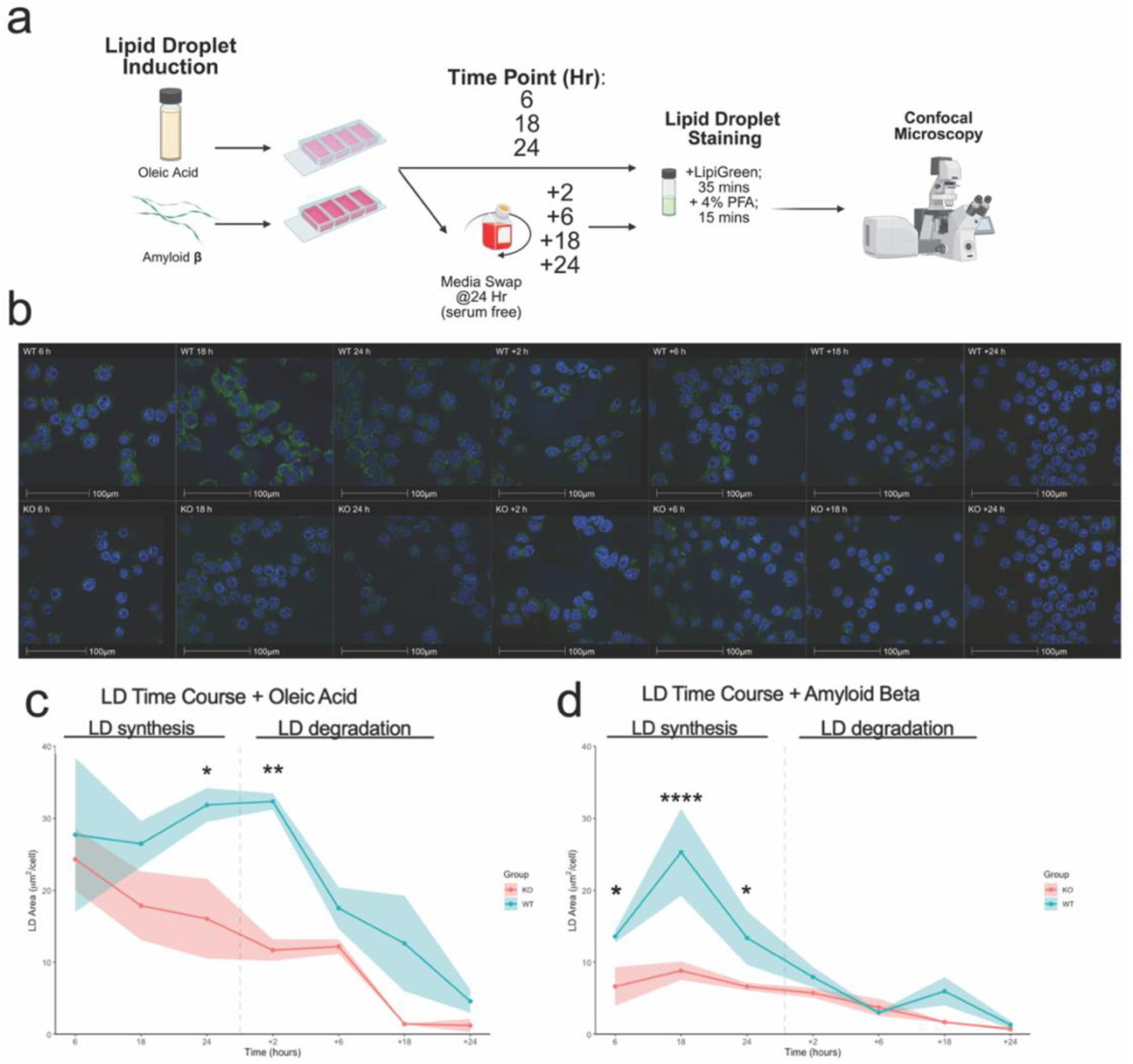
Plin2 loss alters lipid droplet accumulation and dynamics. **a**, Experimental workflow. BV2 cells were exposed to oleic acid (OA) or amyloid beta (Aβ) to induce lipid droplets, fixed at 6, 18, and 24 hours, or switched to serum free medium for a chase and collected/imaged at the indicated times after removal. Cells were stained with LipiGreen for neutral lipids and DAPI for nuclei and imaged by confocal microscopy. **b**, Representative fields for WT and Plin2 KO under Aβ treatment at 6, 18, 24, +2, +6, +18, and +24 hours. LipiGreen signal marks droplets shown in green, DAPI stained nuclei shown in blue. **c**, Time course with OA. Total lipid droplet area per nucleus is plotted over time for WT and KO. WT shows a larger rise at 24 hours which remained +2 hours after serum removal, whereas KO remains lower and returns toward baseline. Shading denotes mean with error as SEM. **d**, Time course with Aβ. WT displays an early rise that peaks by 18 hours and remains above KO at 24 hours, while KO shows no significant increase over the same window. Both genotypes return to baseline by 48 hours.

### Mitochondrial respiration in BV2 microglia under basal and amyloid-β challenge

Given the mitochondrial gene expression differences and unique Aβ response, we next assessed bioenergetics in Plin2 KO microglia. KO cells exhibited significantly reduced basal respiration under both control (**p < 0.01) and Aβ-treated (*p < 0.05) conditions (Fig. 6c). Despite this lower baseline, KO cells maintained the ability to reach similar or higher levels of maximal respiration (Fig. 6d), resulting in a greater spare respiratory capacity (*p < 0.05). Proton leak was consistently lower at baseline and after Aβ exposure (***p < 0.001, ****p < 0.0001; Fig. 6e), reflecting tighter coupling efficiency. For glycolysis, WT cells showed a trend toward higher basal ECAR, and upon Aβ exposure exhibited a significant increase, whereas this elevation in glycolysis was blunted in KO cells (p < 0.05; Fig. 6f). Integration of ECAR and OCR data (Fig. 6g) shows that Plin2 loss establishes a lower energetic set-point – both basal OCR and ECAR are reduced – while preserving respiratory capacity. This profile potentially reflects a more efficient baseline with an enhanced ability to meet sudden energetic demands. In contrast, WT microglia allocate toward higher baseline expenditure and mobilize glycolysis in response to Aβ. Together, these findings suggest that Plin2 functions as a calibrator of microglial energy budgeting: promoting higher baseline activity and glycolytic responsiveness under Aβ treatment.

**Figure 6.**
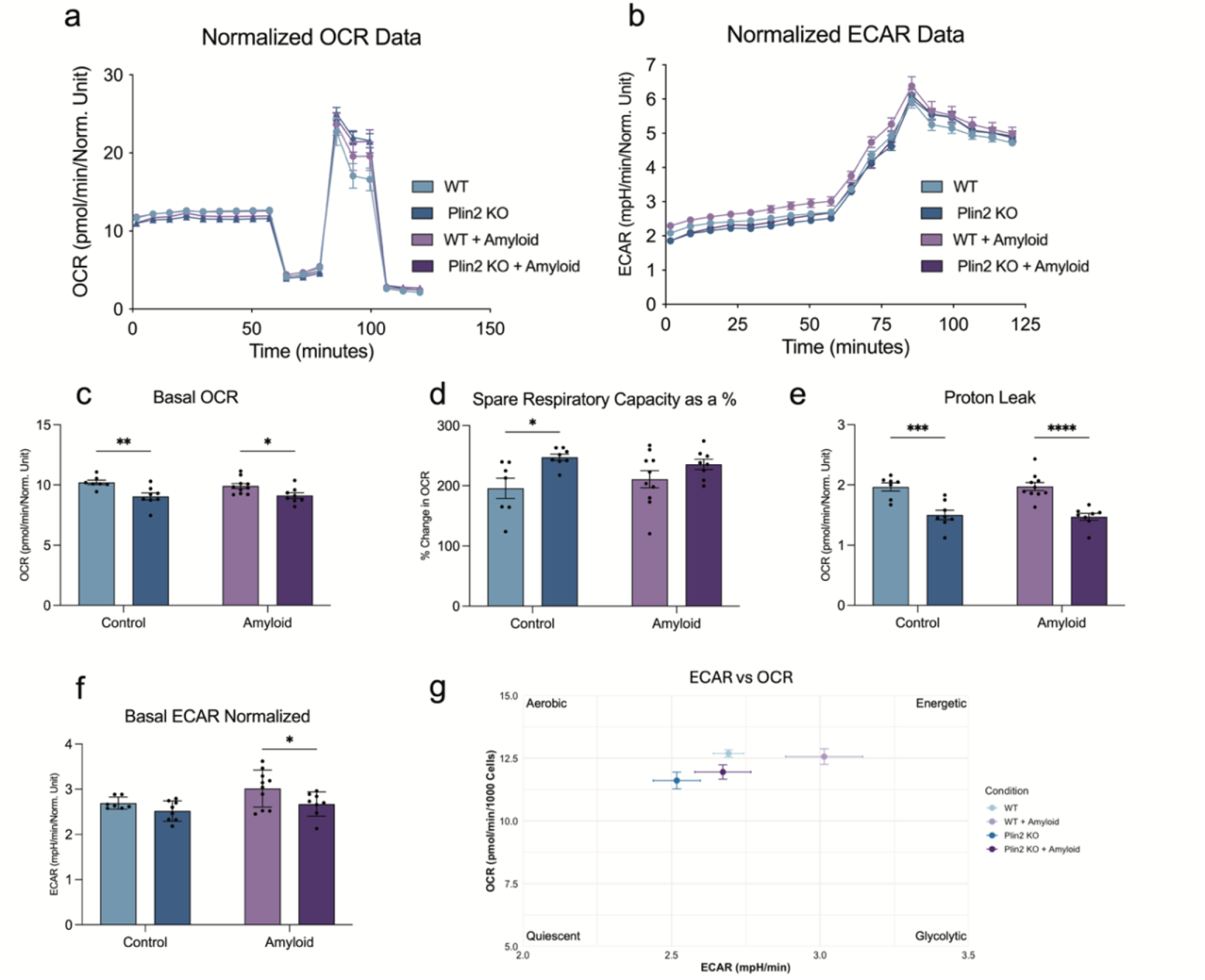
Plin2 deletion reshapes microglial bioenergetics. **a**, Oxygen consumption rate (OCR) traces for WT and Plin2 KO BV2 cells under control or amyloid beta (Aβ) conditions. **b**, Extracellular acidification rate (ECAR) traces for the same groups. **c**, Basal OCR. Plin2 KO operates at a lower basal respiration than WT in both conditions. **d**, Spare respiratory capacity expressed as percentage, calculated from FCCP-stimulated OCR relative to basal and maximal respiration. KO shows a larger reserve. **e**, Proton leak OCR following oligomycin. KO exhibits reduced leak compared to WT. **f**, Basal ECAR. KO relies less on glycolysis under Aβ. **g**, energetic phenotype plot mapping basal ECAR against basal OCR for each group.

### Plin2 KO Significantly Alters the Microglia Lipidome

Because Plin2 regulates neutral lipid storage, we next performed targeted lipidomics to define how its loss reprograms the BV2 lipidome. Across conditions, including control, oleic acid, and dN2A, WT cells consistently accumulated higher levels of triacylglycerols and diacylglycerols, while Plin2 KO cells showed relative enrichment of cholesteryl esters (Fig. 7b–g). Under myelin treatment, however, CE levels did not differ between genotypes, though KO cells continued to display reduced TAG/DAG abundance (Fig. 7g). In our Aβ treated samples, we see a similar trend as before where the KO cells have significantly lower TAG/DAG abundance with an increase in abundance of CE. Previous studies have shown that LD composition is primarily composed of TAG (∼90%)^30^. Thus, the significantly lower abundance of TAG/DAG seen in the KO samples is likely the driving force for the lower LD accumulation. These data support the conclusion that Plin2 promotes TAG-rich droplet storage in microglia, while its loss reduces neutral lipid accumulation and modulates cholesterol handling across diverse stimuli in vitro.

**Figure 7.**
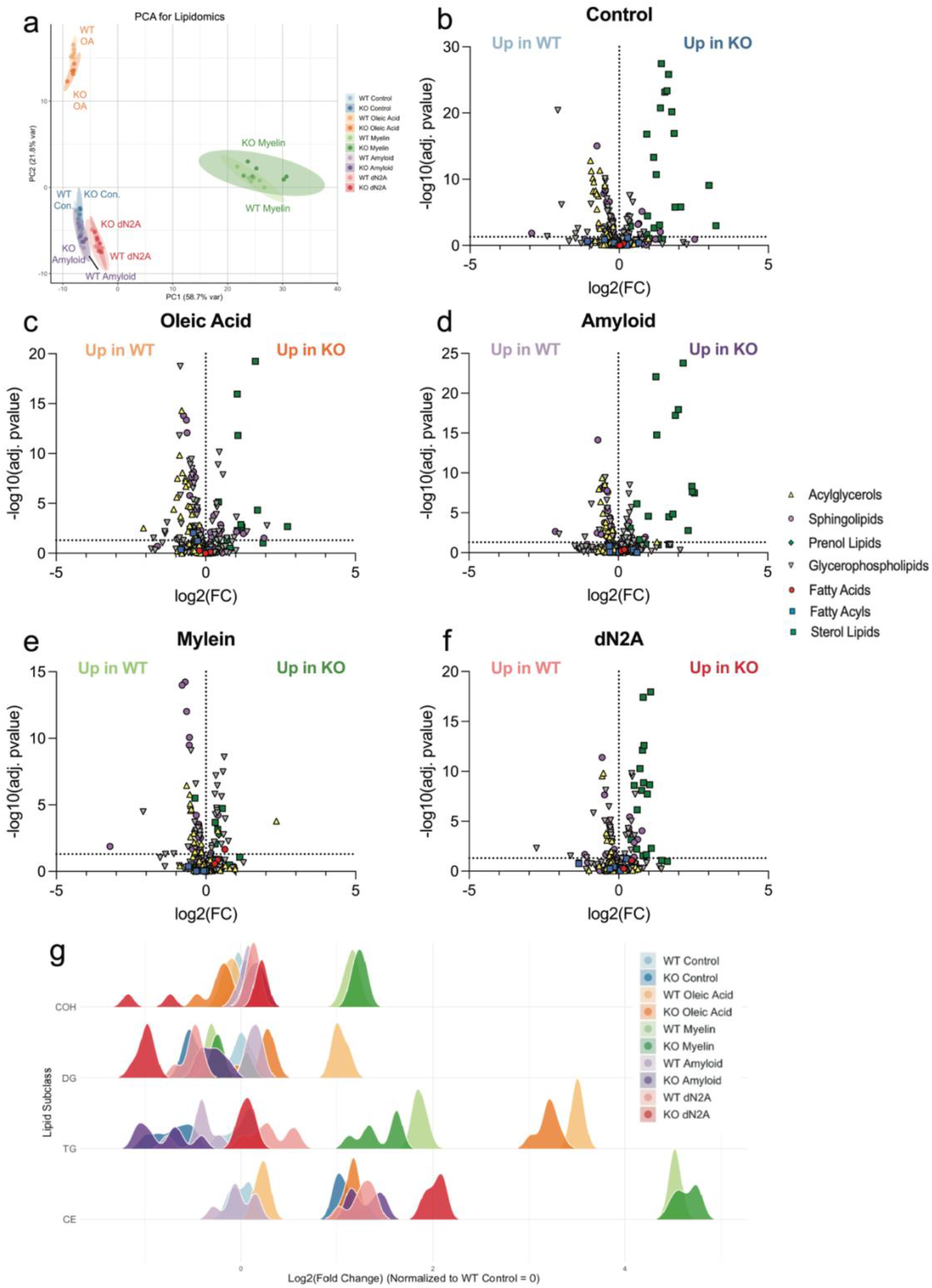
Targeted lipidomics reveals Plin2-dependent routing of neutral lipids. **a**, Principal component analysis of global lipid profiles from WT and Plin2 KO BV2 cells under control, oleic acid (OA), amyloid beta (Aβ), myelin, and dN2A treatments. **b–f**, Volcano plots by condition showing log2 fold change (KO vs WT) on the x-axis and −log10(adjusted p value) on the y-axis. Each point is a lipid species, with shape indicating lipid class (acylglycerols, sphingolipids, glycerophospholipids, fatty acids, prenol lipids, sterol lipids). Labels at top highlight species increased in WT or increased in KO. Across Control, OA, Aβ, and dN2A, WT is enriched for triacylglycerols and diacylglycerols, whereas KO shows higher cholesteryl ester. **g**, Class-level distributions of log2 fold change (KO vs WT) for representative lipid classes across conditions. Density ridges summarize species-level shifts and recapitulate the pattern that WT favors TAG/DAG while KO favors CE.

## Discussion

Microglia play a central role in maintaining brain homeostasis through surveillance, clearance, and immune regulation. A growing body of work demonstrates that LD accumulation fundamentally alters these functions^12,27,31,32^. LD-rich microglia, emerge in aging and AD, where they exhibit impaired phagocytosis, heightened oxidative stress, and pro-inflammatory bias^12,15,31,33^. Despite this recognition, the molecular scaffolds that stabilize LDs in microglia remain poorly understood. Our study identifies the LD coat protein Plin2 as a key determinant of this phenotype, showing that its loss limits droplet expansion and reprograms microglial function under diverse metabolic and disease-relevant challenges. Specifically, we show loss of Plin2 lowers LD accumulation through reduction of TAG abundance, improves phagocytosis capacity, and yields a unique transcriptional and metabolic profile in response to Aβ in microglia.

Work in macrophages and hepatocytes has established Plin2 as a structural protein that shields lipid droplets from lipolysis^23,24,34,35^, thereby promoting triglyceride retention. Our results support this paradigm in microglia. Under oleic acid exposure, WT microglia primarily expanded droplets by increasing size, whereas Plin2 KO cells maintained smaller droplets. This pattern indicates a lesser role for Plin2 in controlling droplet initiation but a rather critical role in stabilizes their growth. Time-course analyses reinforced this conclusion: while WT cells progressively accumulated LDs with oleic acid and exhibited a transient burst under Aβ treatment, KO cells remained comparatively resistant to expansion. Together, these findings support a model in which Plin2 acts as a molecular brake on LD turnover, sustaining accumulation in conditions of lipid excess or amyloid stress.

Lipid-laden phagocytes are consistently characterized by reduced clearance capacity, and defective phagocytosis is a hallmark of AD microglia^12,31^. Our results demonstrate that Plin2 KO enhances uptake of zymosan particles under both basal and lipid-loaded conditions. Zymosan engages CLEC7A/TLR2 and Fc receptors^36^, providing a broad assay of innate uptake capacity. The improved clearance in KO cells mirrors transcriptomic signatures enriched for Fc receptor signaling, innate immune pathways, and actin remodeling, all of which support recognition and internalization. Importantly, inflammatory effectors such as CCL chemokines and matrix metalloproteinases were lower in KO cells, suggesting that loss of Plin2 decouples lipid droplet accumulation from pro-inflammatory output. These observations align with the broader literature in which LD-rich microglia in AD exhibit compromised clearance and inflammatory bias but add a novel mechanistic insight: Plin2 may contribute directly to this dysfunction by stabilizing the droplets that restrain immune adaptability.

Bulk RNA-seq revealed that Plin2 knockout consistently drove down sterol and cholesterol biosynthetic pathways while upregulating programs for chemotaxis, migration, RNA/ribonucleoprotein metabolism, and mitochondrial organization. Importantly, under amyloid challenge, KO cells mounted a distinctive transcriptional response characterized by enrichment of mitochondrial respiratory chain assembly, oxidative phosphorylation, and RNA splicing. This suggests that loss of Plin2 does more than blunt lipid storage, it also enables a transcriptional program that preserves mitochondrial integrity and RNA metabolism in the face of Aβ stress. In the context of AD, where LD-rich microglia are marked by oxidative stress, defective energy production, and impaired clearance^5,12,37^, the KO amyloid signature represents a potentially protective adaptation that contrasts sharply with the lipid burdened state.

WGCNA reinforced this shift at the network level. Down in KO were modules for sterol/cholesterol biosynthesis and TNF superfamily cytokine production, features that mirror the maladaptive, pro-inflammatory lipid burdened signature in aging and AD. By contrast, up in KO modules included Fcγ receptor signaling, innate immune pathways, and Arp2/3-mediated actin remodeling, networks central to receptor-driven phagocytosis, cytoskeletal remodeling, and debris clearance. These adaptive modules parallel those described in microglia that retain phagocytic competence under demyelination and amyloid challenge^14,38^. Thus, Plin2 loss not only silences sterol- and cytokine-driven rigidity but also reconfigures network hubs toward immune engagement and cytoskeletal flexibility, placing Plin2 at the nexus of LD stability, microglial clearance, and AD pathology.

These network changes were borne out in cellular energetics. WT cells mounted strong glycolytic responses to Aβ, consistent with literature showing that inflammatory activation depends on aerobic glycolysis via the mTOR–HIF1α axis^39^. Yet this shift, while acutely supportive of cytokine production, is maladaptive with chronic exposure, ultimately driving collapse of both glycolysis and OXPHOS^40^. Plin2 KO cells blunted this glycolytic surge (lower ECAR) while maintaining maximal respiration and spare capacity. This “low-glycolysis, high-reserve” profile could reflect a form of metabolic resilience, where mitochondrial flexibility is preserved rather than sacrificed for short-term inflammatory gain. Such a state may enable sustained phagocytic surveillance under prolonged amyloid stress, a functional contrast to the exhausted lipid laden phenotype. Further work is needed to fully elucidate these effects under both acute and chronic inflammatory responses.

Targeted lipidomics further showed the significance of removing Plin2. Across control, oleic acid, Aβ, and dN2A cells, WT were enriched for TG and DG, whereas Plin2 consistently showed higher levels of CEs. The simplest interpretation is that Plin2 loss shifts neutral lipid routing away from TG storage and promotes its utilization. This is consistent with much of the previous work showing that loss of Plin2 enhances the turnover of LDs largely through lipolysis^24,41^, and likely through the process of lipophagy^19^. Interestingly, these results match a previous report where fasted Plin2 KO mice had substantial reductions in TG species with modest increases in CEs^42^. However, when mice from this same study were given a high fat diet Plin2 KO still resulted in reduced TGs where CEs were unchanged. Another study has specifically addressed the effect of removing Plin2 on cholesterol toxicity in bone marrow derived macrophages and found it to be well tolerated, like our current study^43^. This evidence suggests the blunted LD accumulation seen in response to Plin2 ablation is largely driven by a reduction in acylglycerols, though increased efflux of cholesterol from Plin2 KO macrophages as also been reported ^24,44^. This suggest that Plin2 also plays a significant role in cholesterol handling in microglia which is of particular interest in neurodegenerative disorders associated with loss of myelination. Full understanding on Plin2’s role in these processes may require further investigation in vivo.

We believe our data support the growing trend of targeting LDs for their role in AD, and that interventions which reduce maladaptive droplet accumulation can improve microglial performance^24,31^. Lipid-laden microglia are a hallmark of the aged brain and are observed near plaques and in tauopathy, where droplet-laden microglia show impaired clearance and pro-inflammatory signatures^12,15,18,33^. This phenotype is further amplified by genetic risk: the strongest common risk factor for late-onset AD, APOE4, is linked to lipid droplet rich microglial states^45^. Our data suggest that loss of Plin2 acts as a lever on this axis, pushing microglia away from chronic lipid storage and toward a more clearance-competent, metabolically resilient state. Thus, we view Plin2 as a novel therapeutic target for lowering lipid laden microglia in AD, one that does not inhibit droplet formation but instead promotes rapid turnover and utilization.

Our study has limitations, including several inherent to the BV2 model. As a transformed cell line, it cannot capture the full heterogeneity, cell-cell interactions, and chronic exposures that shape primary murine or human microglia in vivo^46-48^. Next steps should prioritize validation in primary or induced pluripotent stem cell derived microglia under chronic lipid stress, direct measurement of cholesterol efflux kinetics, and lipophagy flux in the Plin2 knockout background, as well as testing central nervous system restricted Plin2 modulation in models of demyelination and amyloid and/or tau pathology.

In conclusion, this work establishes Plin2 as a central regulator of microglial lipid droplet biology, demonstrating that its loss reshapes transcriptional, metabolic, and functional programs. Removing Plin2 prevents microglia from becoming lipid-burdened, a state that in aging and AD is linked to impaired phagocytosis, chronic inflammation, and metabolic collapse. Instead, Plin2 knockout drives microglia toward a putatively adaptive profile. Therapeutically, modulating Plin2 or droplet turnover could offer a strategy to restore microglial performance in neurodegeneration. This approach is unique in that it lowers the lipid burden while still retaining the ability to generate LDs for lipid buffering and regulation. Together, our data define a novel mechanism by which droplet stability controls microglial fate and provide a foundation for targeting lipid-burdened microglia in disease.

## Supporting information

WGCNA Supplemental Table

## Declarations

## Ethics approval and consent to participate

Not applicable. This study did not involve human participants. All animal work was performed in accordance with institutional guidelines and approved by the University of Kentucky IACUC (2016-2569).

## Consent for publication

Not applicable.

## Availability of data and materials

All data generated or analyzed during this study are included in this published article and its supplementary information files.

## Competing interests

The authors declare that they have no competing interests.

## Funding

This work was supported by the National Institute on Aging R01AG081421 and R01AG080589 (LAJ), the National Institute of General Medical Sciences P20 GM148326 (LAJ), and the Alzheimer’s Association (LAJ).

## Authors’ contributions

IOS designed and performed experiments, analyzed data, and wrote the manuscript. LAJ oversaw the project, provided conceptual input, and edited and wrote the manuscript. Both authors read and approved the final manuscript.

## Acknowledgements

We thank the University of Kentucky Light Microscopy Core, the CNS COBRE Metabolomics Core, and the Markey Cancer Center Redox Metabolism Core for providing resources and expertise throughout this project.

## Notes

### Competing Interest Statement

The authors have declared no competing interest.

